# Changes in the urinary proteome in rats with regular swimming exercise

**DOI:** 10.1101/2021.05.18.444417

**Authors:** Wenshu Meng, Dan Xu, Yunchen Meng, Weinan Zhang, Zhiping Zhen, Youhe Gao

## Abstract

**Purpose:** Urine can sensitively reflect early pathophysiological changes in the body. The purpose of this study was to determine whether the urine proteome could reflect changes in regular swimming exercise.

**Methods:** In this study, experimental rats were subjected to daily moderate-intensity swimming exercise for 7 weeks. Urine samples were collected at weeks 2, 5, and 7 and were analyzed by using liquid chromatography coupled with tandem mass spectrometry (LC-MS/MS).

**Results:** Unsupervised clustering analysis of all urinary proteins identified at week 2 showed that the swimming group was distinctively different from the control group. Compared to the control group, a total of 112, 61 and 44 differential proteins were identified in the swimming group at weeks 2, 5 and 7, respectively. Randomized grouping statistical analysis showed that more than 85% of the differential proteins identified in this study were caused by swimming exercise rather than random allocation. According to the Human Protein Atlas, the differential proteins that have human orthologs were strongly expressed in the liver, kidney and intestine. Functional annotation analysis revealed that these differential proteins were involved in glucose metabolism and immunity-related pathways.

**Conclusion:** Our results revealed that the urinary proteome could reflect significant changes following regular swimming exercise. These findings may suggest an approach to monitoring whether the amount of exercise is appropriate.

## Introduction

Urine is a good source for biomarker discovery. Without homeostatic mechanisms, urine can sensitively reflect early pathophysiological changes in the body, and these changes might be useful disease biomarkers [1]. Since the composition of urine is affected by various factors, such as age, sex, and diet [2–5], animal models are an effective tool to minimize external influencing factors due to their similar genetic backgrounds and the same living environment. Thus, disease animal models can establish relationships between the disease and the corresponding changes in the urine proteome. Our laboratory found that changes in urinary proteins occurred before pathologic or clinical manifestations appeared in various types of animal models, such as subcutaneous tumor model [6], Alzheimer’s disease model [7], chronic pancreatitis model [8], liver fibrosis model [9], and myocarditis model [10]. Recent studies have also shown that the urine proteome has potential for differential diagnosis. For example, early urinary proteins were different when the same tumor cells were grown in different organs [6, 11–14] and when different cells were injected into the same organ [12, 15]. Furthermore, several clinical studies performed urine proteomics to discover diagnostic biomarkers, such as for gastric cancer [16] and familial Parkinson’s disease [17]. However, it is unknown whether urine can show changes in response to regular physical exercise.

Physical exercise as a pathophysiological process that can improve health conditions and has a positive role in the prevention, management, and regulation of numerous chronic conditions [18–20], including cancer and coronary heart diseases[21, 22]. Many studies have shown that exercise has a profound effect on the immune system. Furthermore, it has been demonstrated that physical exercise exerts a positive impact on the nervous system, learning and memory[23–25]. Swimming is a popular physical activity and an effective option for maintaining and improving cardiovascular health. Recent studies have shown that swimming is beneficial for mental health and cognitive ability [26, 27]. To the best of our knowledge, there is no previous study on urine proteomics changes during regular swimming exercise.

Rats have the innate ability to swim and are the first choice for swimming models [28]. In this study, experimental rats were subjected to daily moderate-intensity swimming exercise for 40 min per day for 7 weeks. Urine samples were collected at weeks 2, 5, and 7. The urine proteome was analyzed by liquid chromatography coupled with tandem mass spectrometry (LC-MS/MS). The purpose of this study was to determine whether the urine proteome could reflect changes in regular swimming exercise. The experimental design and workflow of the proteomics analysis in this study are shown in Figure 1.

**Figure 1.**
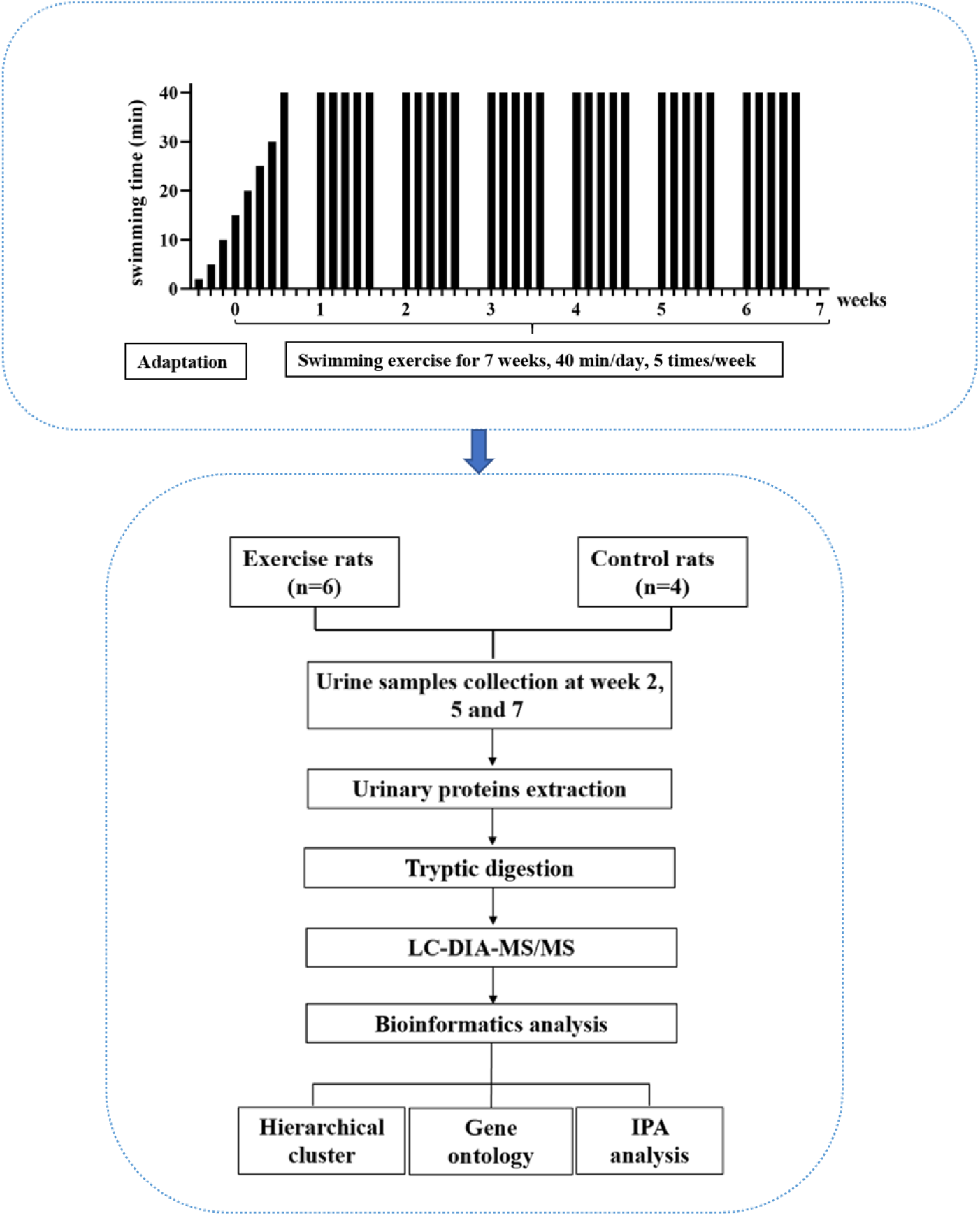
The experimental design and workflow of the proteomics analysis in this study. The experimental rats were subjected to daily moderate-intensity swimming exercise for 7 weeks. Urine samples were collected at week 2, 5, and 7 during swimming exercise. Urine proteins were identified by liquid chromatography coupled with tandem mass spectrometry (LC-MS/MS).

## Materials and methods

### Experimental animals

Male SD rats (seven days old) were supplied by the Department of Neurobiology, School of Basic Medical Sciences, Peking University. All animals were housed with free access to a standard laboratory diet and water with a 12-h light-dark cycle under standard conditions (indoor temperature 22 ± 1°C, humidity 65-70%). The experiment was approved by the Institute of Basic Medical College (ID: ACUC-A02-2014-007). The study was performed according to guidelines developed by the Institutional Animal Care and Use Committee of Peking Medical College.

### Swimming exercise

A large pool (diameter: 1500 mm, height: 500 mm) served as the swimming pool. The water temperature was maintained at 36°C. For the adaptation phase, rats swam for increasing amounts of time, from 2 minutes to 10 minutes over three days. For the exercise phase, the intensity of exercise in the first week gradually increased from 15 minutes to 40 minutes, and the intensity at 40 minutes lasted for 6 weeks, which is considered to be moderate exercise [18]. The animals were quickly and gently dried after each training session. The rats (n=10) were randomly divided into the following two groups: experimental rats (n=6) and control rats (n=4). In the experimental group, the rats underwent the 7-week swimming exercise program. The control rats did not swim.

### Urine collection and sample preparation

Urine samples were collected from the experimental and control groups at weeks 2, 5 and 7 during the swimming exercise. The animals were individually placed in metabolic cages for 10 h to collect urine samples without any treatment. After collection, the urine samples were stored at -80°C. The urine samples (n=30) were centrifuged at 12,000 g for 40 min at 4°C to remove cell debris. The supernatants were precipitated with three volumes of ethanol at -20°C overnight and then centrifuged at 12,000 g for 30 min. Then, lysis buffer (8 mol/L urea, 2 mol/L thiourea, 50 mmol/L Tris, and 25 mmol/L DTT) was used to dissolve the pellets. The protein concentration of the urine samples was measured by the Bradford assay.

### Tryptic digestion

Urinary proteins (100 μg of each sample) were digested with trypsin (Trypsin Gold, Mass Spec Grade, Promega, Fitchburg, WI, USA) using filter-aided sample preparation (FASP) methods [29]. These peptide mixtures were desalted using Oasis HLB cartridges (Waters, Milford, MA) and dried by vacuum evaporation (Thermo Fisher Scientific, Bremen, Germany). The digested peptides (n=30) were redissolved in 0.1% formic acid to a concentration of 0.5 μg/μL. The iRT reagent (Biognosys, Switzerland) was spiked at a concentration of 1:10 v/v into all samples for calibration of the retention time of the extracted peptide peaks. For analysis, 1 μg of peptides from an individual sample was analyzed by LC-DIA-MS/MS.

### Reversed-phase fractionation spin column separation

A total of 90 μg of pooled peptides was generated from 6 μl from each sample and then separated by a high-pH reversed-phase peptide fractionation kit (Thermo Pierce, Waltham, MA, USA) according to the manufacturer’s instructions. A step gradient of increasing acetonitrile concentrations (5, 7.5, 10, 12.5, 15, 17.5, 20 and 50%) was applied to the columns to elute the peptides. Ten different fractionated samples (including the flow-through fraction, wash fraction, and eight step gradient sample fractions) were collected and dried by vacuum evaporation. The ten fractions were resuspended in 20 μl of 0.1% formic acid, and 1 μg of each of the fractions was analyzed by LC-DDA-MS/MS.

### LC-MS/MS analysis

Peptide samples were analyzed in an EASY-nLC 1200 chromatography system (Thermo Fisher Scientific) and an Orbitrap Fusion Lumos Tribrid mass spectrometer (Thermo Fisher Scientific). The samples were loaded onto a trapping column (75 μm × 2 cm, 3 μm, C18, 100 Å) and separated by a reverse-phase analysis column (50 μm × 15 cm, 2 μm, C18, 100 Å). The eluted gradient was 4%-35% buffer B (0.1% formic acid in 80% acetonitrile) at a flow rate of 300 nL/min for 90 min.

To generate the spectral library, 1 μg of each of ten fractions was analyzed in DDA mode. The parameters were set as follows: the full scan ranged from 350 to 1500 m/z with a resolution of 120,000; MS/MS scans were acquired with a resolution of 30,000; the cycle time was set to 3 s; the HCD energy was set to 30%; the autogain control (AGC) target was set to 4e5; and the maximum injection time was set to 50 ms. In DIA mode, 1 μg of each sample was analyzed. The variable isolation window of the DIA method with 36 windows was set (Table S1). The parameters were set as follows: the full scan ranged from 350 to 1500 m/z with a resolution of 60,000; the DIA scan was acquired from 200 to 2000 m/z with a resolution of 30,000; the HCD energy was set to 32%; the AGC target was set to 1e6; and the maximum injection time was set to 100 ms. A quality control sample of a mixture from each sample was analyzed after every six samples.

### Data analysis

The DDA data of ten fractions were searched against the Swiss-Prot rat database (released in 2017, including 7992 sequences) appended with the iRT peptide sequence using Proteome Discoverer software (version 2.1, Thermo Scientific). The search parameters were set as follows: two missed trypsin cleavage sites were allowed; the parent ion mass tolerances were set to 10 ppm; the fragment ion mass tolerances were set to 0.02 Da; the carbamidomethyl of cysteine was set as a fixed modification; and the oxidation of methionine was set as a variable modification. The false discovery rate (FDR) of the proteins was less than 1%. The search results were used to set the DIA method. The DDA raw files were processed using Spectronaut’s Pulsar database (Biognosys, Switzerland) with the default parameters to generate the spectral library. The DIA raw files were processed using Spectronaut for analysis with the default setting. All of the results were filtered by a Q value cutoff of 0.01 (corresponding to an FDR of 1%), and protein identification was based on one unique peptide. Peptide intensity was calculated by summing the peak areas of their respective fragment ions of MS2, and the protein intensity was calculated by summing the intensities of their respective peptides.

### Statistical analysis

The k-nearest neighbor (K-NN) method was used to fill the missing values of protein abundance [30]. Comparisons between experimental and control groups were performed by one-way ANOVA. The differential proteins at weeks 2, 5 and 7 were screened by the following criteria: fold change ≥ 1.5 or ≤ 0.67; and P < 0.05 by independent sample t-test. Group differences resulting in p < 0.05 were considered statistically significant.

### Functional annotation of the differential proteins

DAVID 6.8 (https://david.ncifcrf.gov/) was used to perform the functional annotation of the differential proteins between the experimental and control groups. The canonical pathways were analyzed with IPA software (Ingenuity Systems, Mountain View, CA, USA).

## Results and Discussion

### Urine proteome changes in the swimming exercise rats

In this study, thirty urine samples from three time points (weeks 2, 5, and 7) from six experimental rats and four control rats were used for LC-DIA-MS/MS quantitation. A total of 698 proteins were identified in all urine samples. A quality control sample of a mixture from each sample was analyzed after every six samples. A total of 518 proteins were identified that had a coefficient of variation (CV) of the QC samples below 30%, and all of the identification and quantification details are listed in Table S2.

Unsupervised clustering analysis of all of proteins identified at three time points was performed (Figure S1). We found that the samples at week 2 were clustered together, indicating that swimming exercise has a great impact on urine after 2 weeks. It is speculated that the clustering effect of the samples at weeks 5 and 7 was poor because the body had adapted to long-term exercise. To further characterize the effects of 2 weeks of swimming exercise, all urinary proteins from 10 urine samples between the two groups at week 2 were analyzed by principal component analysis (PCA). As shown in Figure 2A, the swimming exercise rats were differentiated from the control rats. Meanwhile, unsupervised clustering analysis of all urinary proteins from 10 urine samples between two groups at week 2 was performed. As shown in Figure 2B, the proteomics profiles of the swimming group were distinctively different from those of the control group. These results demonstrated that the urinary proteome changed significantly following swimming exercise.

**Figure 2.**
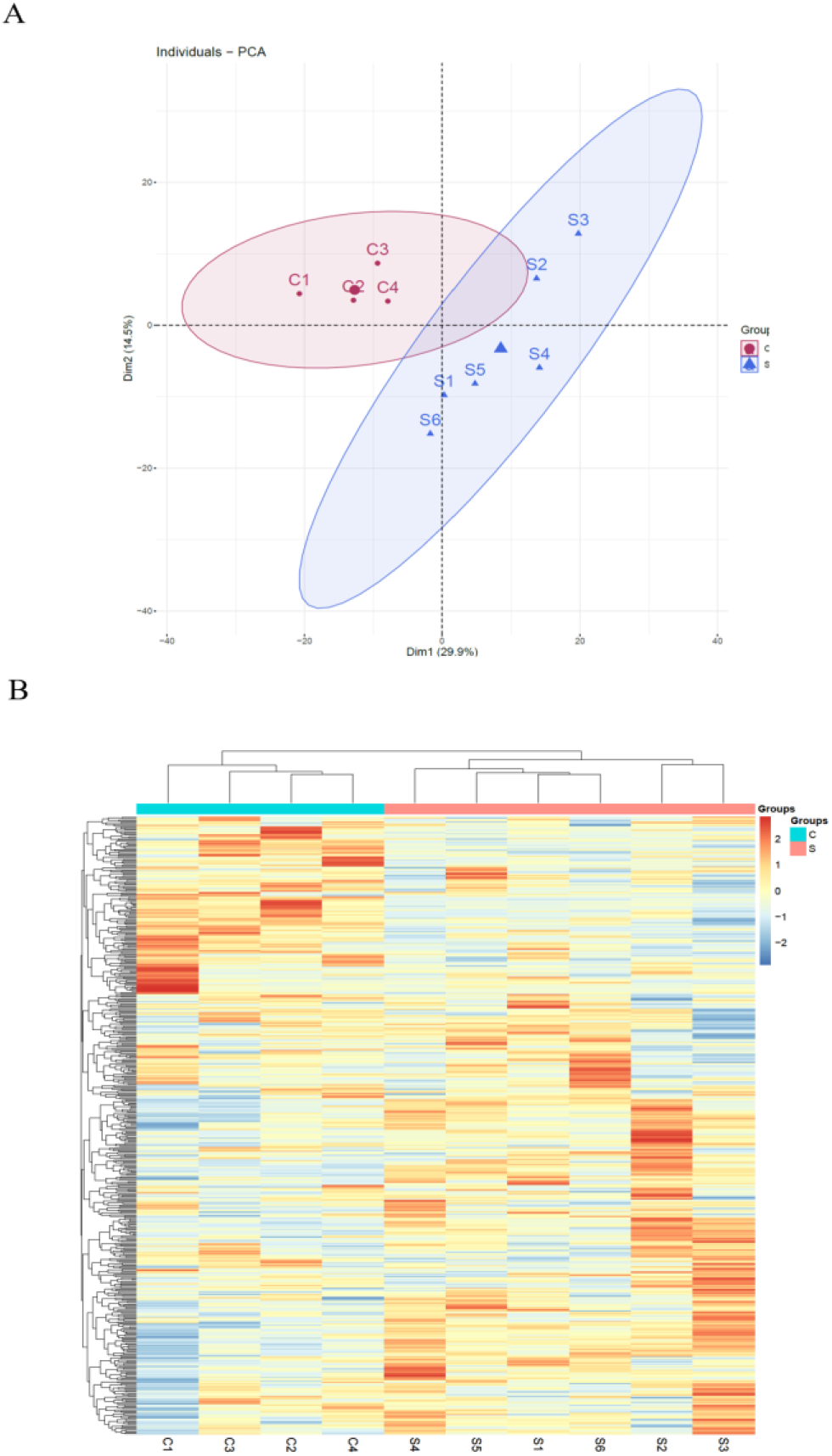
Proteomic analysis of the urine samples of swimming exercise rats. (A) PCA analysis of all proteins from experimental and control urine proteome at week 2. (B) Cluster analysis of all the proteins from experimental and control urine proteome at week 2.

The differential proteins were screened with a *p* value < 0.05 by two-sided, unpaired t-test and a fold change > 1.5 compared with controls. Compared to the control group, 112 differential proteins were identified after 2 weeks of swimming exercise, among which 28 proteins were upregulated and 84 proteins were downregulated (Figure 3A); 61 differential proteins were identified after 5 weeks of swimming exercise, among which 6 proteins were upregulated and 55 proteins were downregulated (Figure 3B); and 44 differential proteins were identified after 7 weeks of swimming exercise, among which 11 proteins were upregulated and 33 proteins were downregulated (Figure 3C). The details of these differential proteins are presented in Table S3. Among these differential proteins, 171 proteins had human orthologs. The overlap of these differential proteins is shown by the Venn diagram in Figure 3D. Five proteins were commonly identified at three time points (Figure 3D), including Ig gamma-1 chain C region, hemopexin, transthyretin, cathepsin D and chondroitin sulfate proteoglycan 4.

**Figure 3.**
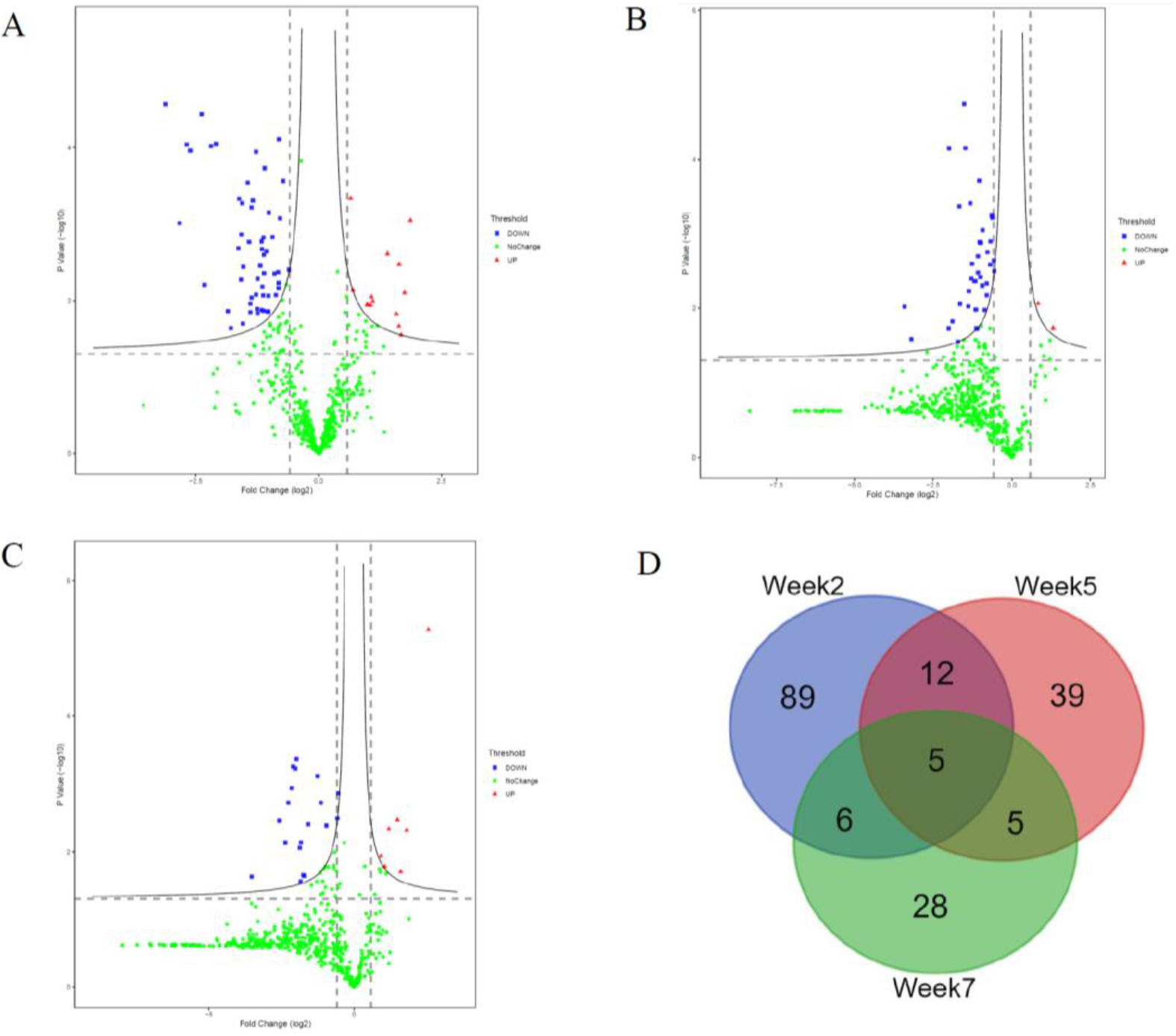
Differential proteins identified between experimental and control group. (A)Volcano plots showing P values (-log_10_) versus protein ratios between experimental and control rats (log_2_) at week 2. (B)Volcano plots showing P values (-log_10_) versus protein ratios between experimental and control rats (log_2_) at week 5. (C)Volcano plots showing P values (-log_10_) versus protein ratios between experimental and control rats (log_2_) at week 7. (D) Overlap evaluation of differential proteins at three time points.

### Tissue distribution of the human orthologs of the differential proteins

To investigate the expression levels of the differential proteins in different tissues and organs, 171 differential proteins that had human orthologs were searched from the Human Protein Atlas. According to the Tissue Atlas, 31 tissues were identified (Figure 4). The differential proteins were strongly expressed in the liver, kidney, intestine, and blood, indicating that these organs may be affected following swimming exercise. Swimming exercise can recruit a large volume of muscle mass. Notably, two proteins, triosephosphate isomerase (TIPSS) and aspartate aminotransferase cytoplasmic (AATC), were strongly expressed in skeletal muscle, indicating that moderate-intensity swimming exercise might have an effect on the muscles of rats.

**Figure 4.**
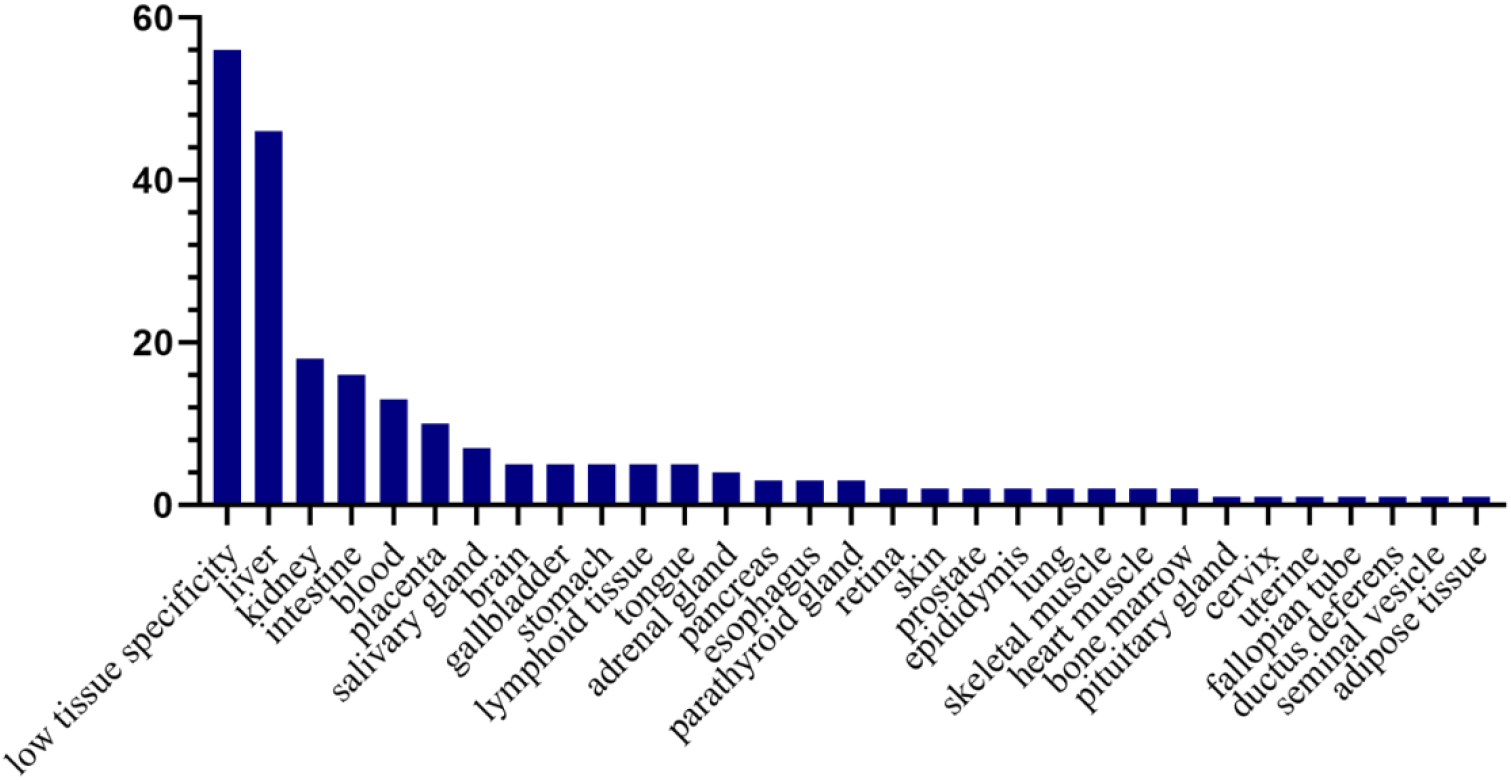
Tissue distribution of the human orthologs of differential proteins. X-axis represents human tissues; Y-axis represent the number of differential proteins.

### Randomized grouping statistical analysis

Considering that omics data are large but the sample size is limited, the differences between the two groups may be randomly generated. To confirm whether the differential proteins were indeed due to swimming exercise, we performed a randomized grouping statistical analysis. We randomly allocated the proteomic data of 10 samples (6 for experimental and 4 for control samples) at each time point and screened for the differential proteins with the same criteria. Then, the average number of differential proteins in all random combinations was calculated, which was the false positive in the actual grouping. There were 210 random allocations at each time point, and the average number of differential proteins in all random combinations at each time point was 15, 5 and 6. The results showed that the false-positive rates were 13.4%, 5% and 13.6% at weeks 2, 5 and 7, respectively. Therefore, most of the differential proteins identified at each time point in this study were caused by swimming exercise rather than random allocation. The details are presented in Table S4. These results suggested that the sample size of this study was sufficient to prove the significant difference in the urine proteome between the swimming group and the control group.

### Functional annotation analysis of the differential proteins

Functional annotation of differential proteins at weeks 2, 5 and 7 was performed by DAVID [31]. The differential proteins identified at three time points were classified into three categories: biological process, cellular component and molecular function.

In the biological process category (Figure 5A), negative regulation of endopeptidase activity and carbohydrate metabolic process were overrepresented at weeks 2 and 5; complement activation, classical pathway, innate immune response and positive regulation of cholesterol esterification were overrepresented at weeks 2 and 7. Response to lipopolysaccharide and positive regulation of cholesterol esterification were only overrepresented at week 2; defense response to bacterium and cell adhesion were only overrepresented at week 5; B cell receptor signaling pathway and positive regulation of B cell activation were only overrepresented at week 7. Notably, some immune-related processes were enriched following swimming exercise. Exercise has a profound effect on immune system function, and studies have shown that regular moderate intensity exercise is beneficial for immunity [32, 33]. Additionally, we found that positive regulation of cholesterol esterification was enriched after swimming exercise in this study. Regular physical exercise provides a wide range of cardiovascular benefits as a nonpharmacological treatment and promotes cholesterol esterification and transport from peripheral tissues to the liver [34, 35].

**Figure 5.**
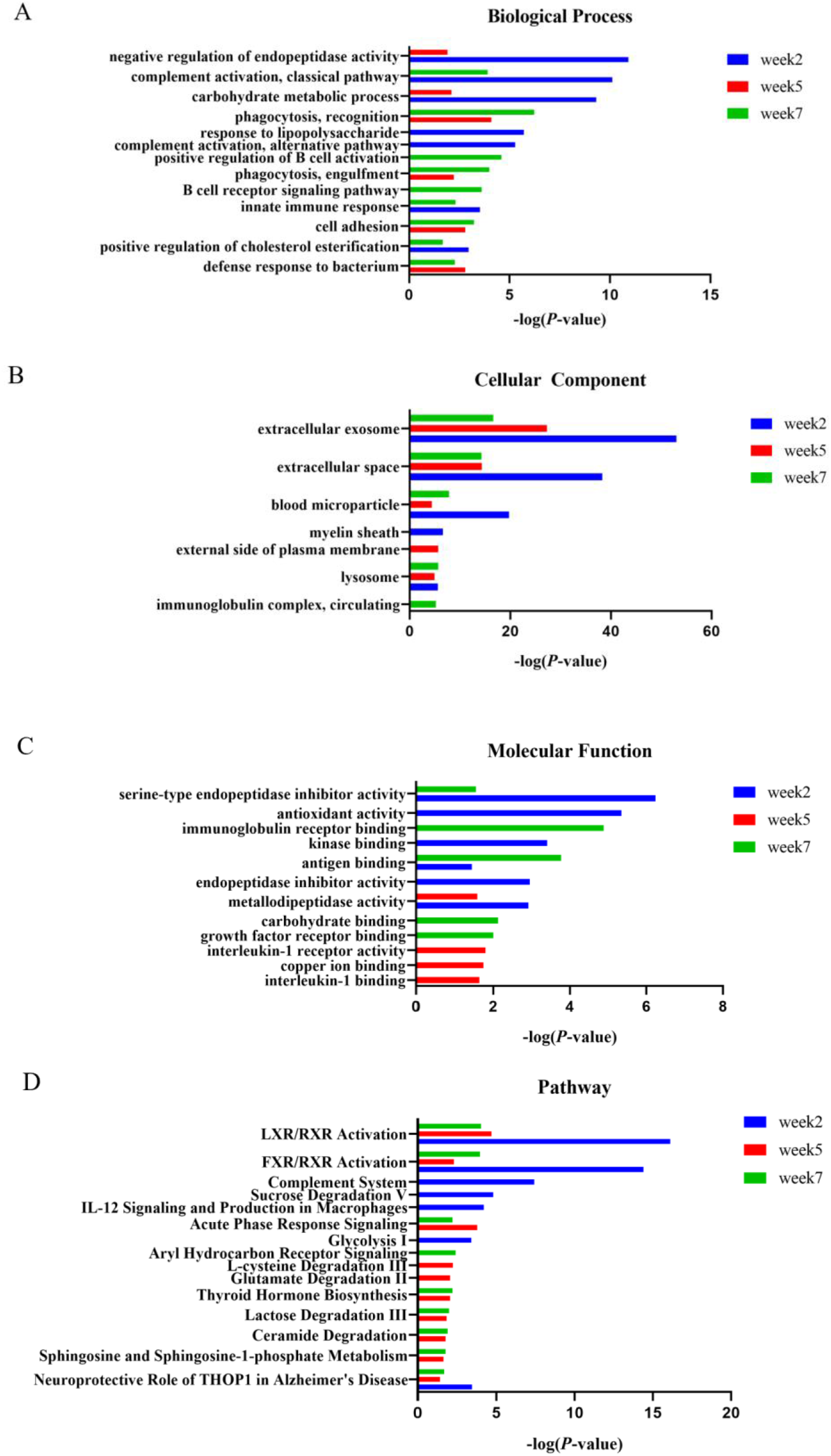
Functional enrichment analysis of differential proteins in this study. (A) Biological process (B) Cellular component (C) Molecular function (D) Canonical pathways

In the cellular component category (Figure 5B), the majority of these differential proteins were from extracellular exosome and extracellular space. In the molecular function category (Figure 5C), metallodipeptidase activity was overrepresented at weeks 2 and 5; antioxidant activity, serine-type endopeptidase inhibitor activity, antigen binding, carbohydrate binding and growth factor receptor binding were overrepresented at weeks 2 and 7.

To characterize the canonical pathways involved with these differential proteins, IPA software was used for analysis. As shown in Figure 5D, LXR/RXR activation and FXR/RXR activation were enriched at three time points. Sphingosine and sphingosine-1-phosphate metabolism, ceramide degradation, lactose degradation III, and thyroid hormone biosynthesis were enriched at weeks 5 and 7. Complement system, sucrose degradation V, IL-12 signaling and production in macrophages, and glycolysis I were enriched at week 2. Glutamate degradation II was enriched at week 5. Aryl hydrocarbon Receptor signaling was enriched at week 7. Among these, some pathways were previously reported to be associated with physical exercise. For example, sphingosine-1-phosphate (S1P) plays an important role in skeletal muscle pathophysiology, and S1P metabolism was found to be regulated by exercise [36]. The S1P content in plasma and its receptors in skeletal muscles were reported to be increased in the skeletal muscle of rats after resistance training [37]. Sphingosine and sphingosine-1-phosphate metabolism were enriched in the urine after swimming exercise in this study. Additionally, carbohydrates are the most efficient fuel for working muscles. The first metabolic pathways of carbohydrate metabolism are skeletal muscle glycogenolysis and glycolysis, and circulating glucose becomes an important energy source. Lactate was also reported to play a primary role as either a direct or indirect energy source for contracting skeletal muscle. We found that some glucose metabolism-related pathways were enriched in urine. Furthermore, glutamate has been implicated in exhaustive or vigorous exercise [38], and a study showed that glutamate increased significantly in the visual cortex following exercise [39]. In this study, we found that glutamate degradation II was enriched in urine following moderate-intensity exercise. Overall, the urine proteome can reflect changes associated with physical exercise.

This study was a preliminary study with a limited number of rats, and the differential proteins identified in this study require further verification in a large number of human urine samples. Urine proteomes after different lengths of exercise were different, suggesting that urine proteomics may distinguish long-term and short-term responses to exercise. Additionally, this is a starting point for further studies of urinary proteome after different types and intensities of exercise to monitor the amount of exercise and to develop an optimal exercise plan. Physical exercise may be an influencing factor in urine proteomics research. When using human urine samples to discover disease biomarkers, physical exercise-related effects can be excluded in future studies.

## Conclusion

Our results revealed that the urinary proteome could reflect significant changes following swimming exercise. These findings may provide an approach to monitoring whether the amount of exercise is appropriate.

## Acknowledgments

This work was supported by the National Key Research and Development Program of China (2018YFC0910202, 2016YFC1306300); the Fundamental Research Funds for the Central Universities (2020KJZX002); Beijing Natural Science Foundation (7172076); Beijing cooperative construction project (110651103); Beijing Normal University (11100704); and Peking Union Medical College Hospital (2016-2.27). The funders had no role in study design, data collection and analysis, decision to publish, or preparation of the manuscript.

## Competing interests

The authors declare that they have no competing interest.

## Supporting information

**Figure S1.**
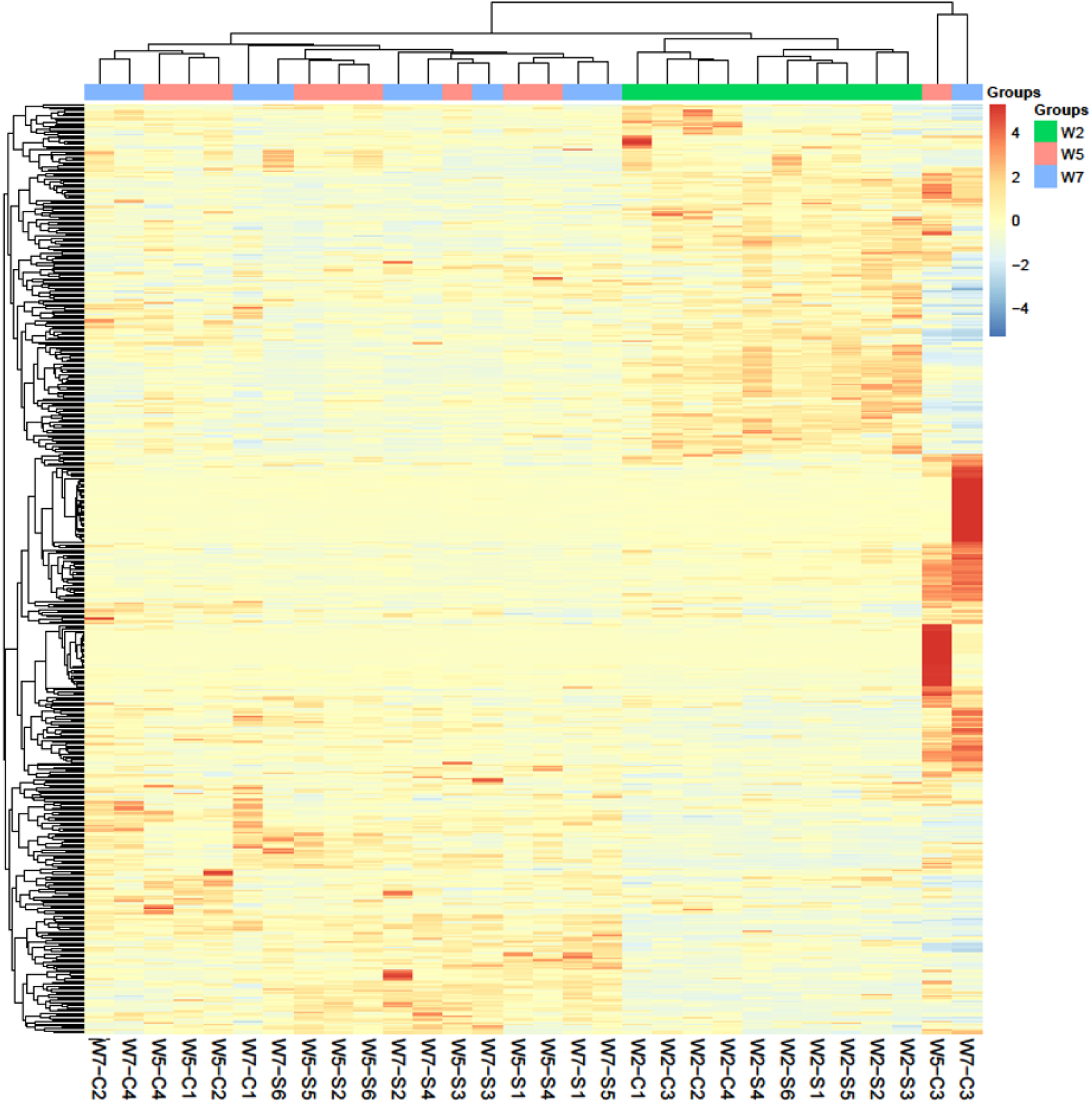
Unsupervised clustering analysis of all of proteins identified at three time points

**Table S1.** The variable isolation window of the DIA method with 36 windows was set for DIA acquisition

**Table S2.** The identification and quantification details of proteins identified in this study

**Table S3.** The details of differential proteins identified at three time points

**Table S4.** The results of randomized grouping statistical analysis

## Notes

### Competing Interest Statement

The authors have declared no competing interest.

